# Balanced RCC1 activity organizes the specialized spindle midplane during cleavage divisions

**DOI:** 10.64898/2026.08.18.745458

**Authors:** Yang Ming, Ai Kiyomitsu, Yutei Takahashi, Tomomi Kiyomitsu

**Affiliations:** Okinawa Institute of Science and Technology Graduate University, 1919-1 Tancha, Onna-son, Kunigami-gun, Okinawa 904-0495, Japan

## Abstract

Chromosome-bound RCC1 generates Ran-GTP signals to organize functional spindles for faithful chromosome segregation during mitosis and meiosis. RCC1 is the sole guanine nucleotide exchange factor (GEF) for Ran and is essential for spindle assembly during early, but not late, embryonic divisions. However, how RCC1 organizes the specialized embryonic spindle and when its function changes during early embryogenesis remain unclear. Here, using time-resolved RCC1 depletion and depletion-rescue experiments in medaka embryos, we show that RCC1 GEF activity is specifically required before the blastula stage to organize a specialized metaphase spindle mid-plane that ensures faithful chromosome segregation. Mechanistically, RCC1 promotes the accumulation of the canonical Ran effectors HURP and KIFC1/HSET, and unexpectedly, the microtubule motor dynein at the spindle midplane during early embryonic divisions. Intriguingly, a five-fold increase in RCC1 expression phenocopies RCC1 depletion, disrupting spindle-midplane organization and the accumulation of KIFC1 and dynein in a GEF activity-dependent manner. Together, our findings demonstrate that both insufficient and excessive RCC1 GEF activity compromise embryonic spindle assembly, revealing that balanced Ran activation is required to organize the specialized spindle midplane during vertebrate cleavage divisions.

**Highlights:**

RCC1 requirement changes with embryonic spindle remodeling before the blastula stage.
RCC1 GEF activity is required to organize the specialized embryonic spindle midplane.
RCC1 promotes the accumulation of HURP, KIFC1, and dynein at the spindle midplane.
Both insufficient and excessive RCC1 GEF activity disrupt the spindle midplane organization.

## Introduction

During mitosis, duplicated chromosomes are captured and segregated by a mitotic spindle, which is composed of numerous short microtubules and microtubule-binding proteins.^1^ Chromosomes are not merely passive cargos but actively generate signals that nucleate and organize spindle microtubules in their vicinity.^2,3^ A central component of chromosome-derived signaling is RCC1 (regulator of chromosome condensation 1), a highly conserved chromatin-binding protein that acts as the sole guanine nucleotide exchange factor (GEF) for the small GTPase Ran.^4,5^ RCC1 generates Ran-GTP, the GTP-bound form of Ran, and promotes nucleocytoplasmic transport during inter-phase and directs spindle assembly and nuclear envelope reformation during and after mitosis, respectively, in animal cells.^6–9^ Ran-GTP activates spindle assembly factors (SAFs) such as HURP and KIFC1/HSET near chromosomes by releasing them from inhibitory importins.^10,11^ The Ran-dependent pathway acts together with the chromosome-derived chromosomal passenger complex (CPC) pathway^12,13^ and centrosome-dependent microtubule nucleation^14^ to efficiently organize bipolar spindles for faithful chromosome segregation^15^, although the relative contribution of each pathway likely varies across developmental stages.^3,15,16^

The Ran-GTP pathway is required for spindle assembly in early, but not late, *Xenopus* egg extracts.^17^ Consistent with this, RCC1 is essential for functional spindle assembly in medaka (*Oryzias latipes*) 4-cell stage embryos, where it promotes formation of a dense microtubule network at metaphase spindle midplane.^18^ In contrast, RCC1 becomes dispensable for spindle assembly and chromosome segregation at the blastula stage, similar to human somatic cells^19^, when spindles lack the dense midplane microtubule network and instead resemble conventional somatic spindles.^18^ These findings point to a transient, developmentally regulated reliance on the Ran-GTP pathway. Yet it remains unclear when RCC1 function changes during early development and whether embryonic spindle assembly requires precisely balanced RCC1 activity.

Medaka embryos are well suited for investigating spindle assembly *in vivo* because they are transparent and undergo planar, synchronous cleavage divisions that enable us to observe spindles in different blastomeres on the same focal plane.^18,20^ The rapid cleavage cycle (approximately 30 min at 26°C) facilitates direct comparison of spindle assembly mechanisms between early and later developmental stages. In addition, mRNA injection, CRISPR-mediated genome editing, and auxin-inducible degron 2 (AID2)-mediated acute RCC1 degradation provide powerful approaches for functional analysis *in vivo*.^18,21^

Here, we investigated the function of RCC1 in medaka embryos by combining live imaging with time-resolved RCC1 depletion and AID2-mediated replacement of RCC1 with GEF-deficient mutants. We show that RCC1 GEF activity is required to organize a dense microtubule network at the metaphase spindle midplane during cleavage divisions. We further show that RCC1 promotes the accumulation of HURP, KIFC1/HSET, and unexpectedly, the microtubule motor dynein at the spindle midplane, thereby organizing a specialized embryonic spindle midplane. Strikingly, RCC1 overexpression phenocopies RCC1 depletion, inducing abnormal spindle assembly and chromosome mis-segregation through its GEF activity. Together, our findings demonstrate that balanced RCC1 GEF activity is essential for organizing the specialized spindle midplane and ensuring faithful chromosome segregation during vertebrate cleavage divisions.

## Results

### Maternal RCC1 drives embryonic divisions until blastula stage

RCC1 localizes to chromosomes after fertilization throughout medaka embryogenesis.^18^ Although zygotic gene activation (ZGA) begins during the early blastula stage (6-8 hours post-fertilization [hpf])^22^, the timing of paternal RCC1 protein expression has remained unclear. To determine this, we crossed females homozygous for RCC1-mCherry (RCC1-mCh) with males homozygous for RCC1-mAID-mClover-3xFLAG (RCC1-mACF) (Fig. 1A) and performed 24-hour live imaging from the 2-cell stage to late neurula stage (Fig. 1B). As expected, maternally derived RCC1-mCh was detected in nuclei throughout embryogenesis. In contrast, paternal RCC1-mACF became detectable only from the early gastrula stage, approximately 3-4 hours after ZGA, and reached a plateau by the mid-gastrula stage (Figs. 1C and 1D, Fig. S1A, and Movie S1). By the late neurula stage, paternal RCC1-mACF accumulated to levels comparable to maternal RCC1-mCh in the small embryonic cells that form the embryo proper (Fig. S1B). In contrast, maternal RCC1-mCh remained more abundant in the larger nuclei of extra-embryonic cells covering the yolk, which emerged between the blastula and gastrula stage (approximately 6-10 hours) (Fig. S1C). These results indicate that maternal RCC1 is the predominant source of RCC1 during embryonic divisions from the one-cell to blastula stage.

**Figure 1.**
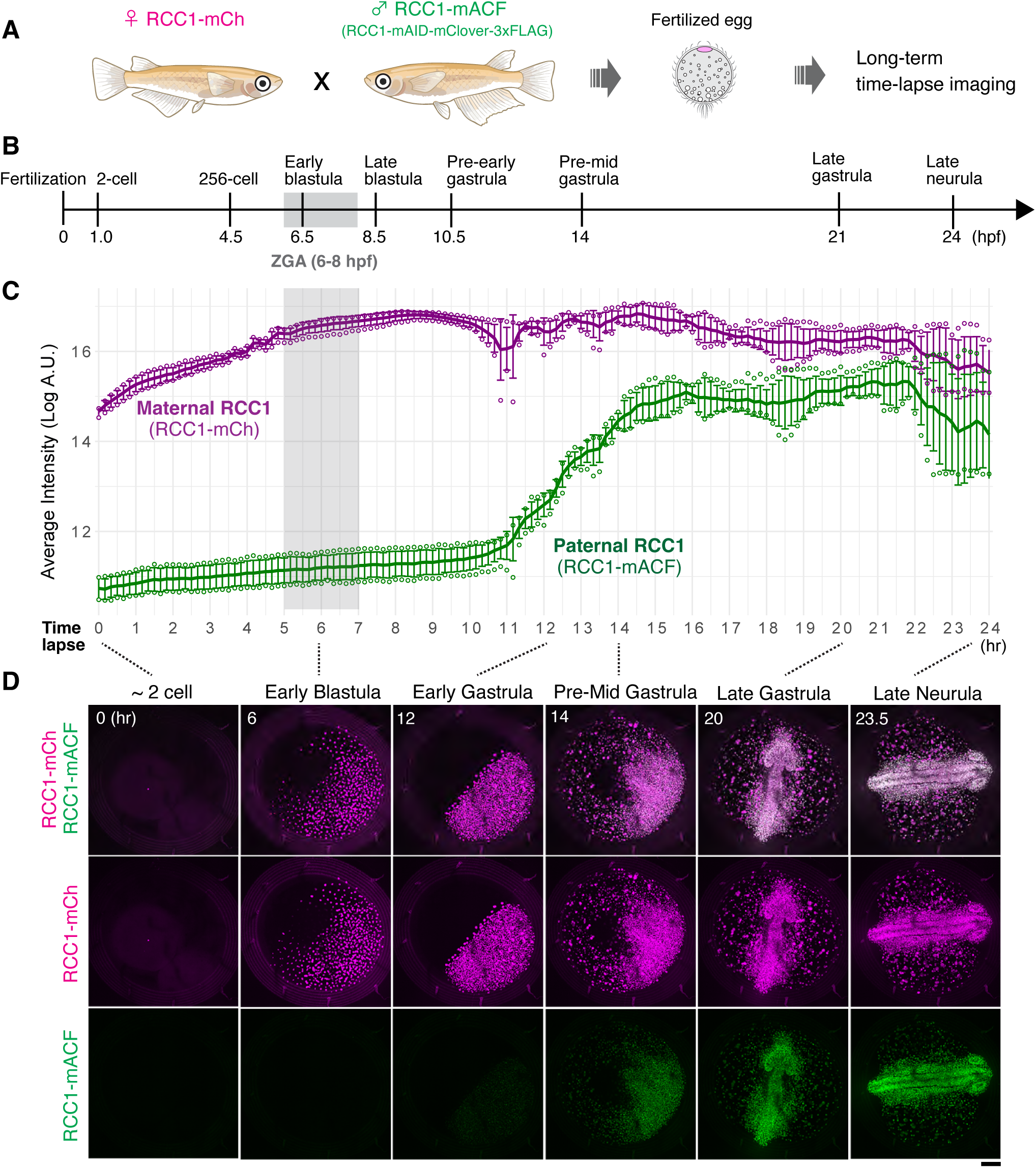
Maternal RCC1 drives embryonic divisions until the blastula stage. (A) Schematic of the experimental design. (B) Schematic showing developmental stages and hours post-fertilization (hpf). (C) Quantification of the mean fluorescence intensities of maternal RCC1 (RCC1-mCh) and paternal RCC1 (RCC1-mACF) during 24-hour time-lapse imaging. Error bars indicate the mean ± SEM and the circles represent the highest and lowest values (n = 5 embryos). (D) Representative live images showing maternal and paternal RCC1 expression from the 2-cell to the late neurula stage. Scale bar, 100 μm.

### RCC1 requirement changes with spindle remodeling during the blastula stage

Embryonic spindle architecture changes dynamically during early medaka development.^18^ In particular, the dense microtubule network at the metaphase spindle midplane, whose formation depends on RCC1, gradually disappears between the 256-cell and early blastula stages^18^ (Fig. 2A and Fig. S2A). To determine when RCC1 is required for spindle assembly and chromosome segregation, we performed time-resolved, AID-mediated RCC1 depletion using the previously established RCC1-mACF medaka strain^18^ (Fig. 2B). Embryos were injected at the one-cell stage with mRNAs encoding OsTIR1(F74G)-P2A-mCherry–Histone H2B (mCh–H2B), followed by the addition of 5-Ph-IAA at 2, 4, 6, or 8 hours post-injection (hpi) to induce RCC1 degradation at defined developmental stages (Figs. 2C and 2D). When RCC1 depletion was initiated at 2 or 4 hpi, nearly all blastomeres exhibited severe chromosome segregation defects (Figs. 2E-2G, Fig. S2B), consistent with our previous observation in 4-cell stage embryos.^18^ In contrast, chromosome segregation defects were rare when RCC1 depletion was initiated at 6 or 8 hpi (Figs. 2H-2L, Fig. S2B). Consistent with these phenotypes, RCC1 depletion abolished the dense microtubule network at the metaphase spindle midplane and markedly altered spindle microtubule intensity profiles at 2 and 4 hpi, but had little effect at 6 hpi (Fig. S2C). Together, these results indicate that RCC1 is required for spindle assembly and faithful chromosome segregation during the first ∼6 hours after fertilization, coinciding with the developmental window in which embryos assemble the specialized spindle architecture.

**Figure 2.**
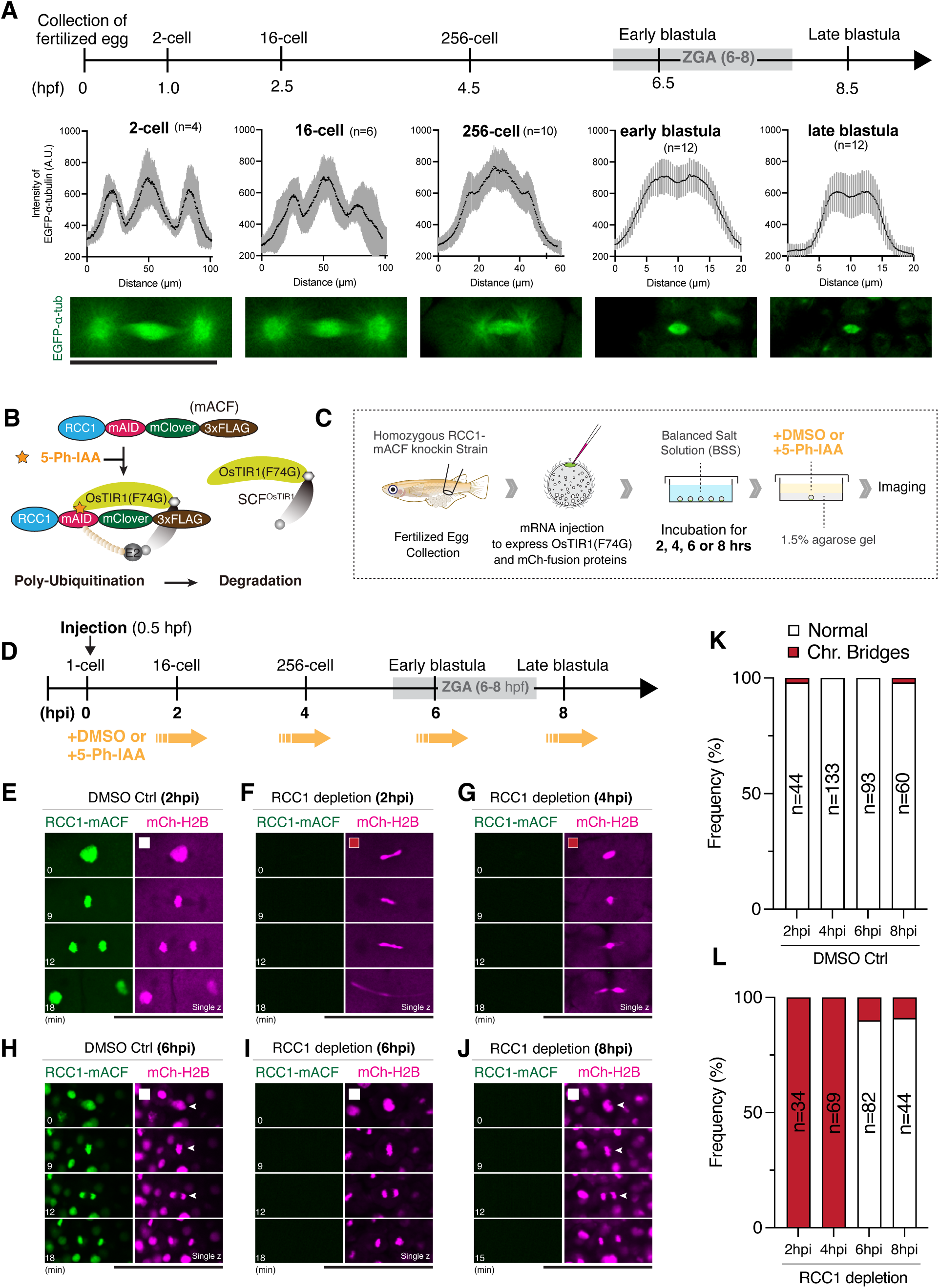
RCC1 requirement changes with spindle remodeling during the blastula stage. (A) Top: Schematic showing developmental stages and hours post-fertilization (hpf). Middle: Line-scan analysis (line widths: 8 px from the 2-cell to early blastula stages; 6 px at the late blastula stage) of EGFP-á-tubulin fluorescence intensity across metaphase spindles showing midplane en-richment before the early blastula stage. Error bars indicate mean ± SD. Bottom: Representative live images of metaphase spindles at the indicated developmental stages. (B) Schematic of auxin-inducible degron 2 (AID2)-mediated RCC1 degradation. (C) Experimental workflow for AID2-mediated RCC1 depletion in fertilized medaka embryos. (D) Schematic of the time-resolved AID2-mediated RCC1 depletion experiment. Embryos were treated with DMSO or 5-Ph-IAA at the indicated time points. (E-J) Representative live images of RCC1-mACF and mCh-H2B in embryos treated with DMSO (control) (E, H) or 5-Ph-IAA (F, G, I, and J) at indicated time points. (K, L) Quantification of chromosome segregation phenotypes in control (K) and RCC1-depleted embryos (L). Scale bars, 100 μm.

### RCC1 GEF activity is required for specialized spindle assembly and chromosome segregation in early embryos

RCC1 is a highly conserved guanine nucleotide exchange factor (GEF) for the small GTPase Ran^5,23^ (Figs. S3A and S3D). To determine which domains of RCC1 are required for embryonic spindle assembly, we performed AID2-mediated protein depletion-rescue experiments using three medaka RCC1 (Ol-RCC1) mutants: RCC1^Δ45^, RCC1^D207A^, and RCC1^D153A/D207A/H324A^ triple mutant (hereafter RCC1^TM^) (Fig. 3A, Fig. S3D). RCC1^Δ45^ lacks the N-terminal nuclear localization signal (NLS) and a medaka-specific 25 aa N-terminal extension (Figs. S3B and S3D). The D207A substitution in RCC1 has been shown to impair GEF activity, whereas RCC1^TM^ carries three substitutions (D153A/D207A/H324A) (Fig. S3C) that are predicted to abolish GEF activity based on studies of human RCC1.^24–26^ AlphaFold 3 modeling further predicts that these conserved residues contribute to the interface with GDP-bound Ran (Fig. 3B).

**Figure 3.**
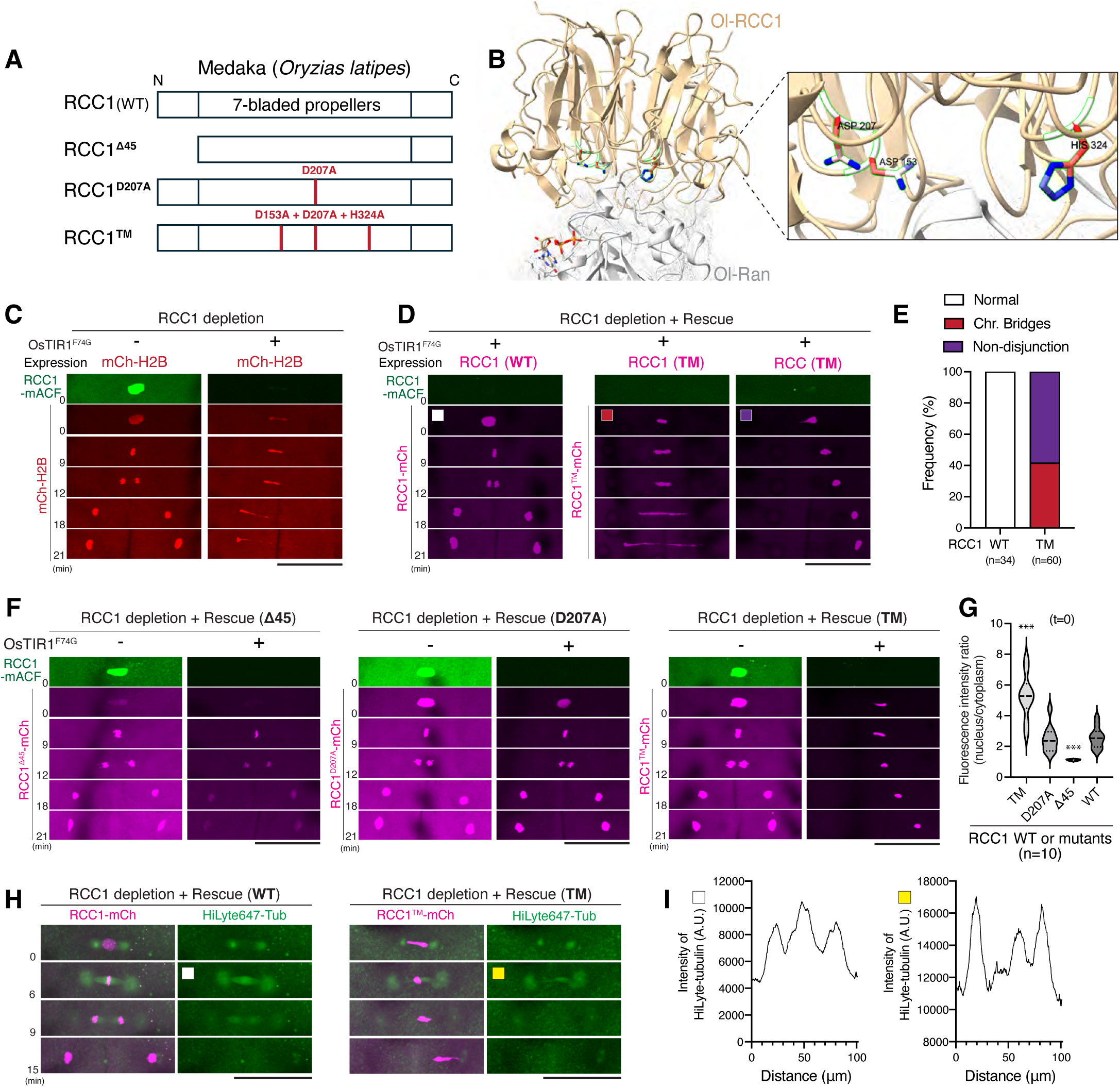
RCC1 GEF activity is required for specialized embryonic spindle assembly and faithful chromosome segregation. (A) Schematic of the RCC1 mutants used in this study. (B) AlphaFold 3-predicted structure of the medaka RCC1-Ran-GDP complex. (C) Representative maximum-intensity-projection (MIP) live images of control (left) and RCC1-depleted (right) 4-cell blastomeres showing chromosome segregation defects following RCC1 depletion. (D) Representative MIP live images showing chromosome segregation phenotypes after replacement of endogenous RCC1-mACF with mCherry-tagged wild-type RCC1 (left) or RCC1^TM^ (right). (E) Quantification of abnormal chromosome segregation in (D). (F) Representative MIP live images of 4-cell blastomeres in which endogenous RCC1-mACF was replaced with mCherry-tagged RCC1^Δ45^ (left), RCC1^D207A^ (middle), or RCC1^TM^ (right). (G) Quantification of the nuclear fluorescence intensity of exogenously expressed mCherry-tagged RCC1 constructs one frame before NEBD in (F, t=0). Statistical significance was assessed using Brown-Forsythe and Welch ANOVA with Dunnett’s multiple comparisons test. \*\*\**p* < 0.001. (H) Representative MIP live images of mitotic spindles in RCC1-depleted 4-cell blastomeres after replacement with wild-type RCC1 (left) or RCC1TM (right). (I) Line-scan analysis (line width 8 px) of HiLyte-tubulin fluorescence intensity along the spindle axis in (H). Scale bars, 100 μm.

To simultaneously degrade endogenous RCC1 and express exogenous RCC1 variants, we injected mRNAs encoding OsTIR1(F74G)-P2A-mCherry-tagged RCC1 constructs into one-cell RCC1-mACF embryos and analyzed 4-cell embryos. Expression of wild-type RCC1-mCherry (RCC1-mCh), but not mCherry-H2B (mCh-H2B), fully rescued the chromosome segregation defects caused by RCC1 depletion (Figs. 3C-3E), confirming the functionality of the rescue assay. In contrast, replacement with RCC1^TM^ failed to rescue chromosome segregation defects despite localizing to mitotic chromosomes similarly to wild-type RCC1 (Figs. 3D and 3E, Movie S2). RCC1^Δ45^ showed reduced nuclear accumulation during interphase, consistent with the loss of its NLS-containing N terminus (Fig. 3F, t=0), but localized normally to mitotic chromosomes and rescued the depletion phenotype (Fig. 3F, Movie S3). Likewise, RCC1^D207A^ localized normally and restored chromosome segregation (Fig. 3F, Movie S4), indicating that the D207A mutation alone is insufficient to disrupt RCC1 function in medaka embryos.

To determine whether GEF activity is required for embryonic spindle architecture, we visualized spindle microtubules by injecting HiLyte 647-tubulin following RCC1 replacement. In contrast to wild-type RCC1, RCC1^TM^ failed to restore the dense microtubule network at the spindle midplane (Figs. 3H and 3I). We also noted that embryos expressing RCC1^TM^ contained smaller, irregularly shaped interphase nuclei before mitotic entry (Figs. 3D and 3H, Fig. S3E), suggesting that RCC1 GEF activity contributes to maintaining nuclear integrity before mitosis. Together, these findings demonstrate that robust RCC1 GEF activity is required for specialized spindle assembly and faithful chromosome segregation in medaka early embryos.

### RCC1 promotes HURP and dynein accumulation at the metaphase spindle midplane

To determine which SAFs are regulated by the RCC1-Ran pathway in early embryos, we analyzed the localization of medaka HURP and KIFC1, two canonical Ran-regulated SAFs^10,11,19^, following mRNA injection in one-cell embryos. As expected, HURP accumulated at the metaphase spindle midplane in both 4-cell (Fig. 4A) and blastula-stage embryos (Fig. 4B), despite the absence of the dense microtubule network at the spindle midplane in blastula-stage embryos. Like HURP, KIFC1 also accumulated around the metaphase spindle midplane in early embryos (Fig. S4A).^27^ Unexpectedly, we found that the microtubule motor dynein, visualized using either DHC-mCh or DHC-mACF^28^, also accumulated at the metaphase spindle midplane in early embryos (Fig. 4C), but not in blastula-stage embryos (Fig. 4D). Long-term live imaging further showed that the dynein accumulation at the metaphase spindle midplane disappears during early blastula stage, coincident with embryonic spindle remodeling (Fig. 4E).

**Figure 4.**
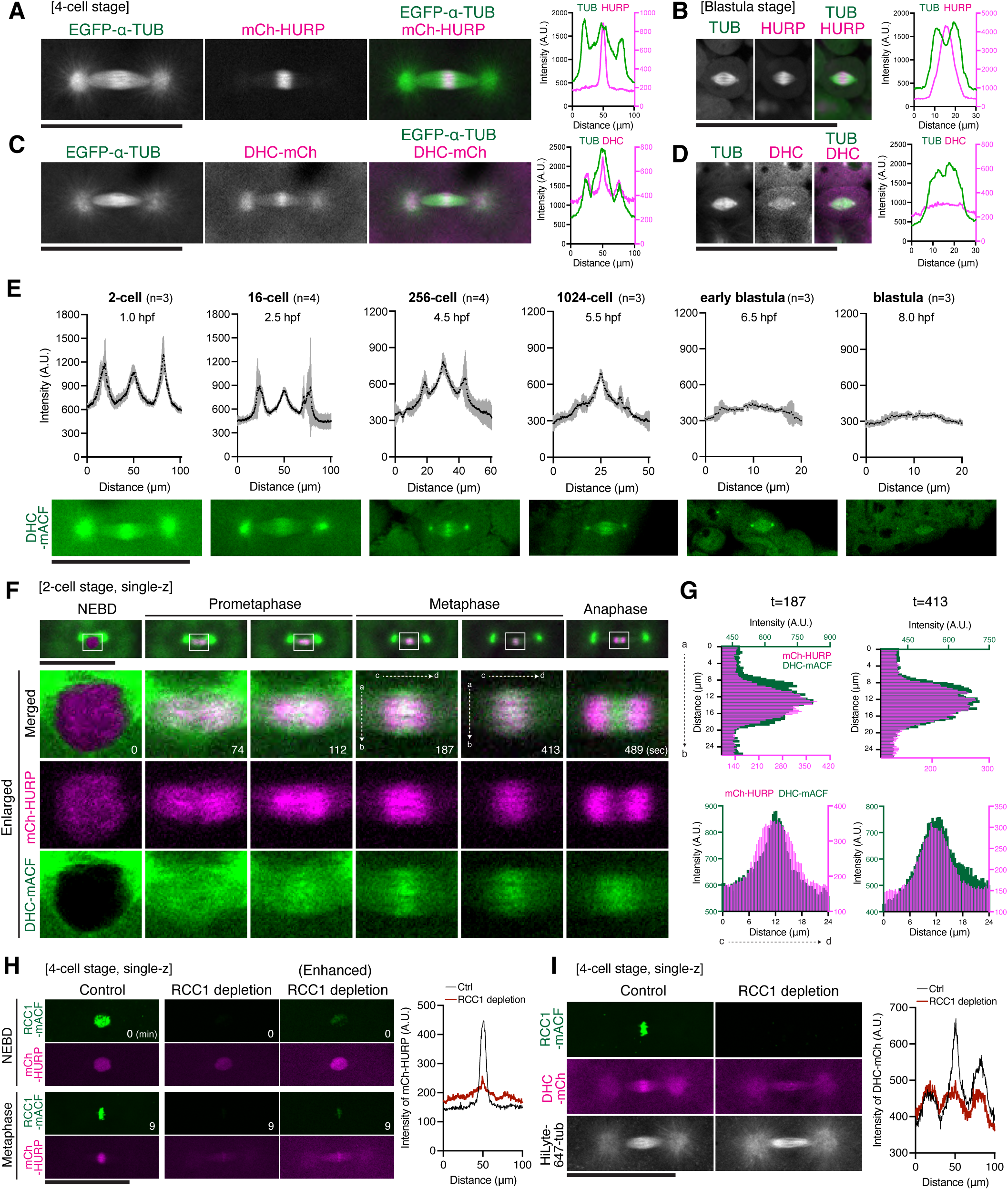
RCC1 promotes HURP and dynein accumulation at the metaphase spindle midplane. (A, B) Representative live images (left) and line-scan analyses (right) showing mCh-HURP accumulation at the metaphase spindle midplane in the 4-cell-stage (A) and blastula-stage (B) embryos (line widths: 8 px in A; 6 px in B). (C, D) Representative live images (left) and line-scan analyses (right) showing DHC-mCh accumulation at the metaphase spindle in 4-cell-stage embryos (C), but not blastula-stage embryos (D) (line widths: 8 px in C; 6 px in D). (E) Top: Line-scan analysis (line widths: 8 px from the 2-cell to 1024-cell stages; 6 px from the early to late blastula stages) of DHC-mACF fluorescence intensity across metaphase spindles showing midplane enrichment before the early blastula stage. Error bars indicate the mean ± SD. Bottom: Representative live images of DHC-mACF localization at metaphase during the indicated developmental stages. (F) Time-lapse images of mCh-HURP and DHC-mACF from NEBD to anaphase. (G) Line-scan analyses (line widths: 16 px for the vertical a-b line; 12 px for the horizontal c-d line) of HURP and dynein fluorescence intensity across the spindle midplane during early (left, t=187) and late metaphase (right, t=413). (H) Representative live images showing reduced nuclear localization of mCh-HURP at NEBD (top) and reduced spindle-midplane accumulation at metaphase (bottom) in RCC1-depleted 4-cell blastomeres. Line-scan analysis (line width 8 px) of mCh-HURP at metaphase is shown on the right. (I) Representative live images (left) and line-scan analysis (right) showing loss of DHC-mCh accumulation at the metaphase spindle midplane in RCC1-depleted 4-cell blastomeres (line width 8 px). Scale bars, 100 μm.

To compare the accumulation dynamics of HURP and dynein at the spindle midplane, we analyzed the localization of HURP, dynein, and microtubules during spindle assembly in 2-cell-stage embryos (Fig. 4F, Fig. S4B, Movie S5). HURP localized to the nucleus in interphase (Fig. 4F), but rapidly associated with spindle microtubules, preferentially in chromosome-proximal regions, after nuclear envelope breakdown (NEBD) (Figs. S4B and S4C). During prometaphase, HURP gradually accumulated at the spindle midplane and showed a single metaphase plate-like localization that separated toward the spindle poles during anaphase, consistent with its localization to kinetochore microtubule plus-ends, as reported in somatic cells.^10^ In contrast, dynein was excluded from the nucleus in interphase but rapidly accumulated throughout the spindle region after NEBD (Fig. 4F). During prometaphase, a subpopulation of dynein accumulated between the HURP-enriched regions in a kinetochore-like pattern. Although this dynein population appeared to dissociate from kinetochores, as observed in somatic cells^29^, a prominent pool of dynein persisted at the spindle midplane throughout metaphase in medaka early embryos (Fig. 4F and Fig. S4B). At early metaphase, dynein was more tightly concentrated at the spindle midplane than HURP, whereas HURP displayed a broader distribution along the spindle axis (Fig. 4F and 4G). As metaphase progressed, the distributions of dynein and HURP increasingly overlapped, becoming largely coincident by late metaphase (Fig. 4F and 4G). During anaphase, dynein localized between the separated HURP signals, suggesting that dynein associates with interpolar rather than kinetochore microtubules.

Importantly, RCC1 depletion markedly reduced the accumulation of exogenously expressed HURP (Fig. 4H, Figs. S4D and S4E), KIFC1^27^, and endogenous dynein (Fig. 4I) at the metaphase spindle midplane. Together, these results demonstrate that RCC1 promotes the accumulation of both the canonical Ran effectors HURP and KIFC1 and, unexpectedly, dynein at the spindle midplane, identifying dynein as a previously unrecognized component of the RCC1-dependent embryonic spindle assembly pathway.

### Embryonic spindle assembly requires balanced RCC1 GEF activity

Since RCC1 GEF activity is required for early embryonic divisions, we next asked whether spindle assembly is sensitive to RCC1 dosage, specifically whether excessive RCC1 activity is also detrimental. We injected RCC1-mACF embryos with high amounts of mRNAs encoding wild-type or mutant RCC1 to induce overexpression (Fig. 3A) and analyzed their phenotypes. Unexpectedly, overexpression of wild-type RCC1, RCC1^Δ45^, or RCC1^D207A^ caused abnormal chromosome segregation (Fig. 5A, Fig. S5A), which was rarely observed when the GEF-deficient RCC1^TM^ mutant was expressed (Fig. 5A). Consistently, overexpression of wild-type RCC1, but not RCC1^TM^ mutant, resulted in embryonic lethality 1 day post fertilization (dpf) (Fig. 5B). To control for variation in expression levels, we compared blastomeres expressing similar levels of mCherry fluorescence in 16-cell-stage embryos. Under these conditions, overexpression of wild-type RCC1, but not H2B or RCC1^TM^, induced lagging chromosomes and chromosome bridges during anaphase (Figs. 5C and 5D). Image quantification showed that cells exhibiting abnormal chromosome segregation had a five- to sixfold higher RCC1-mCh intensity at metaphase chromosomes (Fig. 5E).

**Figure 5.**
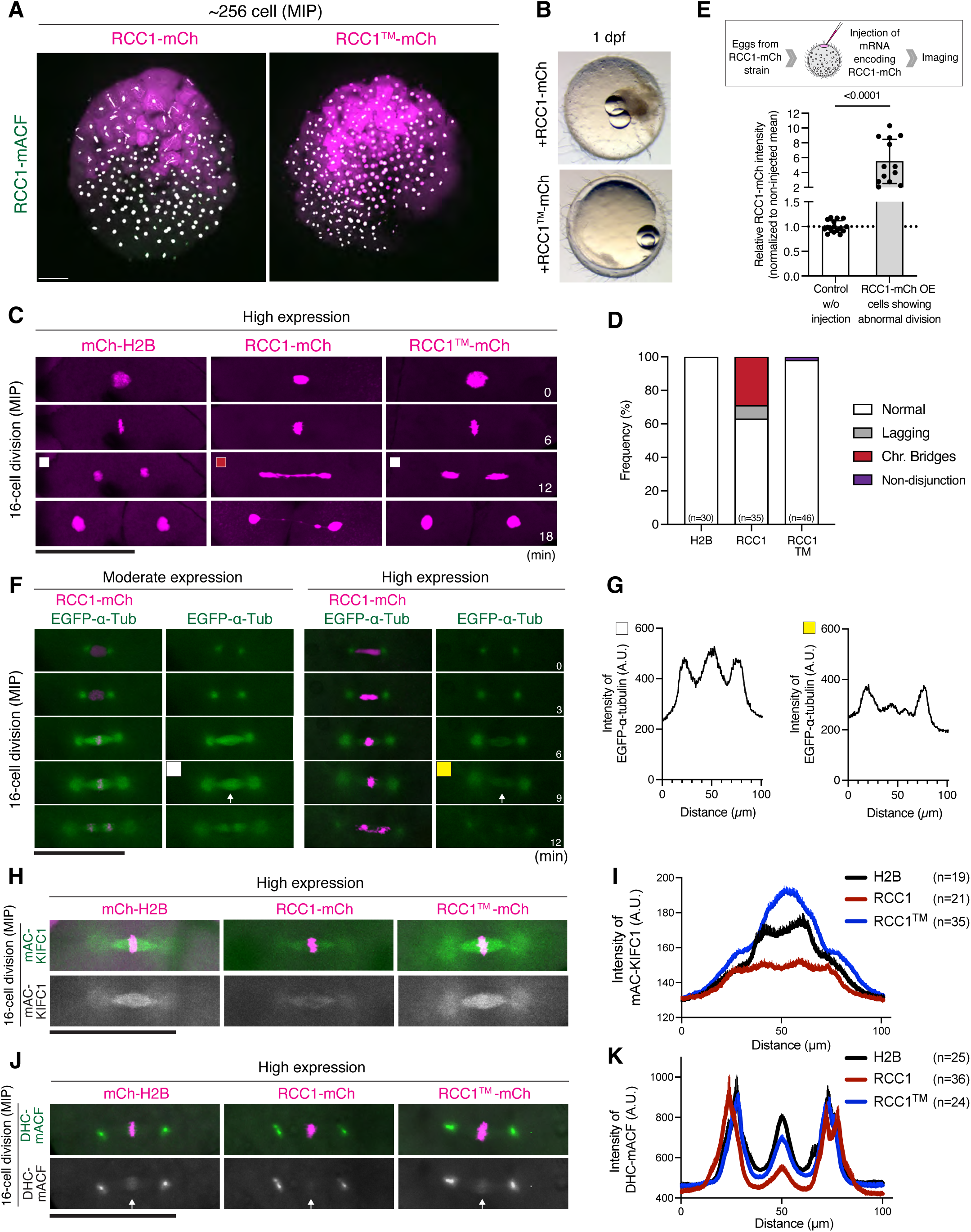
RCC1 overexpression disrupts spindle midplane organization in a GEF activity-dependent manner. (A) Representative live images of 256-cell-stage embryos overexpressing wild type RCC1 (left) or RCC1^TM^ (right). Expression levels varied among blastomeres, likely reflecting uneven distribution of the injected mRNA following injection into one-cell-stage embryos. (B) Phase-contrast images of embryos 1 day after overexpression of RCC1 or RCC1^TM^. Overexpression of wild type RCC1 caused embryonic lethality. (C) Representative live images showing chromosome segregation phenotypes in embryos overexpressing H2B (left), wild type RCC1 (middle), or RCC1^TM^ (right) at the 16-cell stage. (D) Quantification of chromosome segregation phenotypes shown in (C). (E) Quantification of RCC1-mCh fluorescence intensity at metaphase chromosomes in non-injected control and RCC1-mCh-overexpressing blastomeres exhibiting abnormal chromosome segregation. Homozygous RCC1-mCh knock-in embryos were used. Error bars indicate mean ± SD. Two-sided Welch’s t-tests were performed. (F) Representative live images showing metaphase spindle midplanes in blastomeres expressing moderate (left) or high (right) levels of RCC1. (G) Line-scan analysis (line width 8 px) of EGFP-á-tubulin fluorescence intensity across metaphase spindles in (F). (H, J) Representative live images showing the localization of endogenous KIFC1 (mAC-KIFC1) (H) or dynein (DHC-mACF) (J) in embryos overexpressing H2B (left), wild type RCC1 (middle), or RCC1^TM^ (right) at the 16-cell stage. (I, K) Line-scan analyses (line widths: 8 px in I; 10 px in K) of mAC-KIFC1 (I) or DHC-mACF (K) fluorescence intensity across metaphase spindles showing reduced spindle-midplane accumulation of KIFC1 (I) or dynein(K) in RCC1-overexpressing blastomeres. Error bars indicate mean ± SEM. Scale bars, 100 μm.

To determine how excess RCC1 disrupts chromosome segregation, we next visualized microtubules, KIFC1, and dynein following wild-type RCC1 overexpression. Overexpression of RCC1, but not H2B or RCC1^TM^, reduced microtubule density at the spindle midplane (Figs. 5F and 5G, Figs. S5B and S5C). Additionally, RCC1 overexpression abolished the accumulation of endogenous KIFC1 (mAID-mClover-KIFC1) (Figs. 5H and 5I)^27^ and dynein (DHC-mACF)^28^ at the metaphase spindle midplane (Figs. 5J and 5K). Collectively, these findings reveal that both insufficient and excessive RCC1 GEF activity disrupt spindle midplane organization and chromosome segregation, indicating that specialized embryonic spindle assembly requires balanced RCC1 GEF activity.

## Discussion

### RCC1 GEF activity is required to organize the specialized metaphase spindle midplane during cleavage divisions

In this study, we show that maternally supplied RCC1 organizes a specialized, dense microtubule network at the metaphase spindle midplane through its GEF activity in medaka early embryos (Figs. 3D, 3E, 3H, and 3I). RCC1 also promotes the accumulation of the canonical Ran effectors HURP and KIFC1 and, unexpectedly, dynein at the metaphase spindle midplane (Figs. 4H, 4I, and 6A)^27^, suggesting that the RCC1-Ran pathway organizes a specialized spindle midplane that supports faithful chromosome segregation during cleavage divisions.

**Figure 6.**
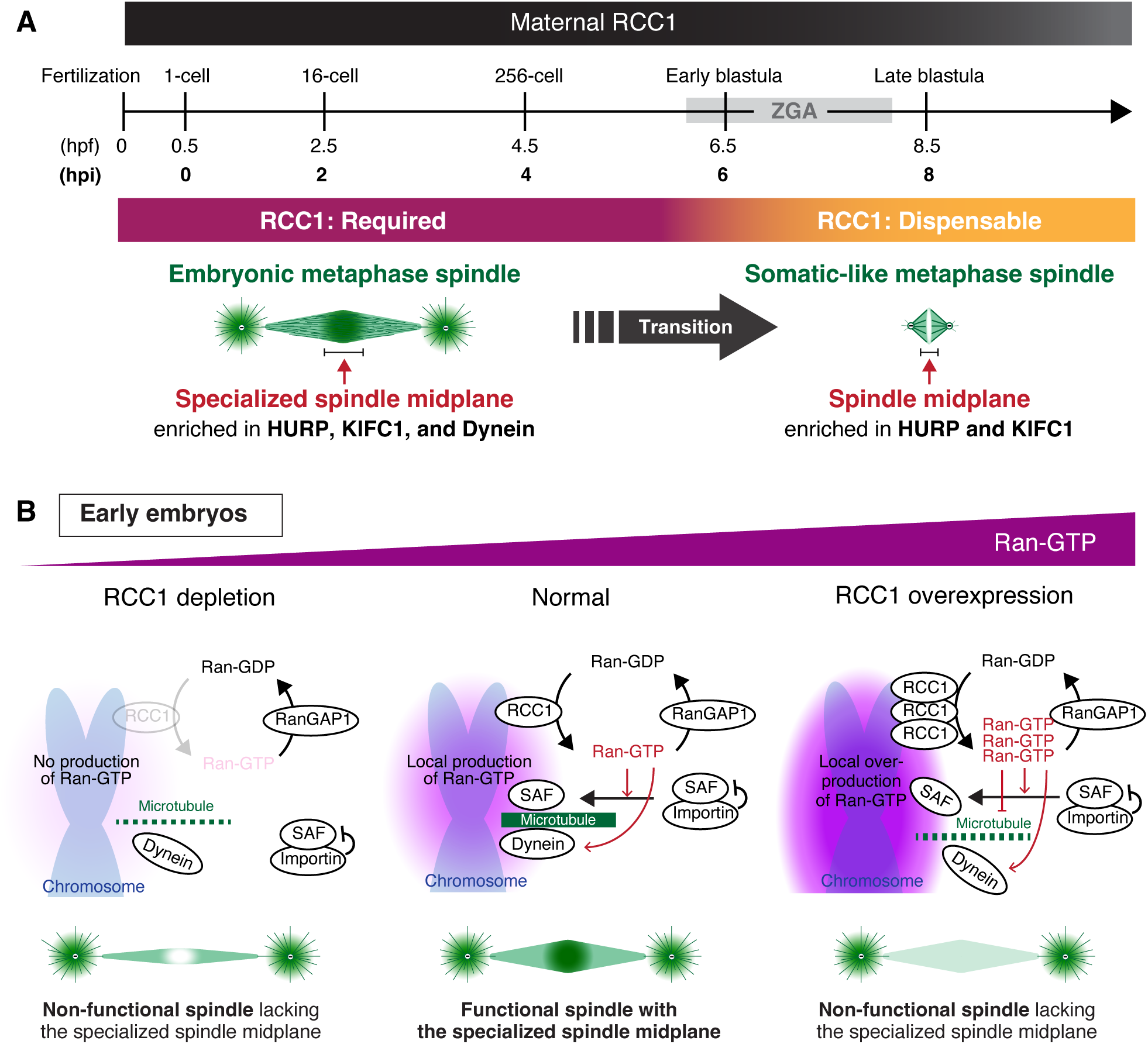
Model illustrating the time- and dosage-dependent contributions of RCC1 to embryonic spindle assembly. (A) Before zygotic genome activation (ZGA), the RCC1-Ran pathway is required to organize the specialized spindle midplane by promoting accumulation of the canonical Ran effectors HURP and KIFC1 while additionally recruiting dynein to the spindle midplane. (B) Both insufficient and excessive RCC1 GEF activity disrupt the specialized metaphase spindle-midplane organization by destabilizing microtubules near chromosomes. Thus, balanced Ran activation is required for embryonic spindle assembly and faithful chromosome segregation during cleavage divisions.

Although Ran-GTP likely activates HURP and KIFC1 by releasing them from inhibitory importins^19,30^, how the RCC1-Ran pathway promotes dynein accumulation at the metaphase spindle midplane remains an important question. Dynein and dynactin localize to the cytoplasm and nuclear envelope during interphase but rapidly accumulate throughout the spindle region after NEBD^28^ (Fig. 4F, Fig. S4B). One possibility is that Ran-GTP promotes dynein recruitment by regulating dynein cofactors, such as LIS1 or Nde1^31–33^, or as-yet-unidentified embryo-specific adaptors immediately before or after NEBD. Our recent study showed that dynein inhibition reduces centrosome-mediated microtubule nucleation in medaka embryos but has only a modest effect on chromosome-mediated microtubule nucleation^28^, suggesting that dynein primarily contributes to spindle microtubule organization rather than chromosome-mediated microtubule nucleation. Because dynein remains associated with interpolar microtubules after anaphase onset, it may stabilize antiparallel microtubules at the spindle midplane during metaphase and at spindle midzone during anaphase^34^ (Fig. 4F). In addition to dynein recruitment, other embryo-specific Ran-dependent mechanisms are likely required to generate the specialized, dense microtubule network at the embryonic spindle midplane. Candidates include the ELYS-Nup107-160 complex^35,36^, which promotes kinetochore-mediated microtubule nucleation, and the augmin-TPX2 pathway^37–45^, which drives microtubule-dependent branching microtubule nucleation.

### Embryonic spindle assembly requires balanced Ran activation

Another key finding of this study is that an approximately fivefold increase in RCC1 protein expression phenocopies RCC1 depletion (Fig. 5). Like RCC1 depletion, RCC1 overexpression destabilizes spindle microtubules around chromosomes and reduces the accumulation of KIFC1 and dynein at the metaphase spindle midplane in a GEF activity-dependent manner (Fig. 5H-K, Fig. 6B). In cultured mammalian cells^46^ and tumors^47^, elevated RCC1 expression is associated with enhanced proliferation, resistance to DNA damage-induced senescence, and poor clinical prognosis, suggesting that increased RCC1 levels are broadly tolerated or even advantageous.^48^ In sharp contrast, excess RCC1 is acutely detrimental to spindle assembly in medaka early embryos, highlighting the distinct requirements for spindle assembly in embryonic and somatic cells. Consistent with this idea, constitutively active RanQ69L and dominant-negative RanT24N induce similar spindle defects during meiosis II^49^, suggesting that specialized embryonic and meiotic spindles are sensitive to both increased and reduced Ran pathway activity. Together, these findings support a model in which specialized embryonic spindle assembly requires precisely balanced Ran activation.

How does excess RCC1 impair specialized embryonic spindle assembly? This effect depends on RCC1 GEF activity, yet expression of constitutively active Ol-RanQ69L rarely caused comparable chromosome segregation defects^18^, arguing against a simple model in which elevated Ran-GTP alone is sufficient to disrupt spindle assembly. Instead, excessive local Ran-GTP production near chromosomes may perturb the finely tuned Ran signaling required for embryonic spindle assembly. One possibility is that locally elevated Ran-GTP promotes excessive GTP consumption, thereby reducing the availability of GTP-bound tubulin for microtubule nucleation and polymerization around chromosomes. Alternatively, active Ran may directly promote microtubule depolymerization near chromosomes, as recently demonstrated *in vitro*.^50^ The gel-like properties of the cytoplasm in large embryos^51,52^ may further amplify these effects by limiting the diffusion of Ran-GTP and GTP around chromosomes. We also found that both RCC1 depletion and overexpression caused abnormal nuclear morphology in a GEF activity-dependent manner (Figs. 3H and 5F), raising the possibility that both reduced and excessive Ran-GTP impair nuclear integrity, thereby indirectly compromising spindle assembly during early embryonic divisions.

In summary, our findings demonstrate that balanced RCC1 GEF activity is required to organize the specialized spindle midplane in medaka early embryos. More broadly, these findings indicate that faithful spindle assembly during vertebrate cleavage divisions depends not simply on the presence of Ran-GTP signaling, but on its precise quantitative regulation.

### Limitations of the study

The medaka RCC1^TM^ mutant was designed based on previous biochemical and structural studies of human RCC1^24–26^ and is predicted to be GEF-deficient, but its GEF activity has not been directly validated *in vitro*. In addition, Ran-GTP gradients were not directly measured in embryos following RCC1 depletion or overexpression. Furthermore, the mechanism by which RCC1 promotes dynein accumulation and the functional role of dynein at the spindle midplane remain unclear. Finally, although medaka embryos provide a powerful vertebrate model for studying embryonic spindle assembly, whether our findings are broadly applicable to other vertebrate embryos remains to be determined.

## Acknowledgments

We thank Toane Arata, Sumika Hagihara, Yoko Nakasone and the OIST animal resource section staffs for feeding and maintenance of medaka fish at OIST. We are grateful to NBRP Medaka (https://shigen.nig.ac.jp/medaka/) for providing OK-Cab (Strain ID: MT830).

## Author contributions

Conceptualization, TK and YM; Investigation, YM, AK, YT; Formal analysis, YM and YT; Methodology, YM, AK and TK; Resources, AK and TK; Writing-original draft, YM; Writing-review & editing, YM and TK; Supervision, TK; Funding Acquisition, TK and AK.

## Data and materials availability

All data supporting findings of this study are available in the paper and its Supplementary Materials. All data of this study are stored at the corresponding author and available on reasonable request.

## Declaration of generative AI and AI-assisted technologies in the writing process

During the preparation of this work, the authors used ChatGPT (OpenAI) to improve the language and readability of the manuscript. After using this tool, the authors reviewed and edited the content as needed and take full responsibility for the content of the published article.

## Methods

### Fish maintenance

All fish experiments were conducted in accordance with protocols (ACUP-2023-009, ACUP-2025-038, ACUP-2025-068) approved by the Animal Care and Use Committee of the Okinawa Institute of Science and Technology Graduate University (OIST). The OK-Cab strain (MT830) of medaka (*Oryzias latipes*) was obtained from the National Bio-Resource Project Medaka (NBRP Medaka) and used as the parental strain. Medaka were raised and maintained as described previously.^18,28^ Naturally fertilized eggs were collected from breeding pairs aged 2–9 months. Only healthy fertilized eggs were selected for imaging. The medaka strains used in this study are listed in Table S2.

### Plasmid construction

To generate constructs expressing EGFP- or mCherry-tagged proteins, the coding sequences (CDSs) of *Oryzias latipes* (Ol) RCC1 (XM_023964257.1), HURP (XM_004080213.5), and KIFC1 (XM_004074065.5) were synthesized by Eurofins (Japan). RCC1 mutant constructs were generated by PCR-based mutagenesis using PrimeSTAR Max DNA Polymerase (Takara). The coding sequence of OsTIR1(F74G)-P2A was amplified from pMK411 (Addgene #140659) and fused in frame with mCh-H2B, mCh-α-tubulin, mCh-RCC1, mCh-HURP, mCh-KIFC1, and mCherry-tagged RCC1 mutants by PCR. All constructs and mutations were verified by Sanger sequencing. The plasmids used in this study are listed in Table S1.

### *In vitro* transcription of mRNA

For *in vitro* transcription, template plasmids were linearized with either NotI or BssHII. mRNAs were synthesized using the mMessage mMachine SP6 Transcription Kit (Thermo Fisher Scientific, AM1340) according to the manufacturer’s instructions. The synthesized mRNAs were purified using the RNeasy Mini Kit (Qiagen).

### Microinjection

Microinjection was performed as described previously.^18^ Glass needles were made from borosilicate glass capillaries using a needle puller (PC-100, Narishige), and mounted on a capillary holder connected to a Femto Jet 4i microinjector (Eppendorf). Needles were manipulated manually using a micromanipulator (MN-153, Narishige) mounted on a stereomicroscope (Leica M80). To express exogenous fluorescent fusion proteins, mRNAs (150 ng/µL) were injected into one-cell-stage embryos. For the co-injection of HiLyte 647-tubulin (Cytoskeleton, Inc., TL670M) and mRNAs, a mixture containing 800 ng/μL HiLyte 647-tubulin and 150 ng/µL mRNA was injected using Sigmacote-treated glass needles (Sigma-Aldrich, SL2-25ML).

### Live Imaging

Live imaging was performed using a spinning-disc confocal microscope equipped with a 20× /0.95 NA water-immersion objective (APO LWD 20× WI λS, Nikon), 488-, 561-, and 640-nm lasers (Coherent), a CSU-W1 spinning-disk confocal unit (Yokogawa Electric Corporation), and an ORCA-Fusion sCMOS camera (Hamamatsu Photonics) mounted on an ECLIPSE Ti2-E inverted microscope with a perfect focus system (Nikon). Immersion water was automatically supplied to the objective using a water-immersion dispenser (Ti2-N-WID, Nikon). For imaging, embryos were mounted in a custom-made agarose chamber prepared in glass-bottom dishes (CELLview™, #627860, Greiner Bio-one). Seven #1.5 coverslips (18 mm×18 mm, 0.12–0.17 mm thickness, Matsunami) were stacked to generate a coverslip mold, which was used to form a shallow concave pocket in the agarose. After solidification, the coverslip mold was removed, producing a pocket approximately 0.8–1.2 mm wide and 3–5 mm deep on the glass surface. Four to six embryos were aligned within the pocket with the blastodisc facing the glass bottom. Dishes were filled with approximately 2 mL of medaka balanced salt solution (BSS, 0.65% NaCl, 0.04% KCl, 0.02% MgSO_4_・7H_2_O, 0.02% CaCl_2_・2H_2_O, sterilized, pH 8.3, adjusted with 5% NaHCO_3_) ^18^, and imaging was performed at room temperature (24-25°C).

### AID2-mediated protein knockdown

AID2-mediated protein degradation was performed as described previously.^18^ One-cell-stage embryos were injected with mRNAs encoding OsTIR1(F74G)-P2A-mCherry fusion proteins. After injection, embryos were gently agitated and cultured in BSS for 10–30 min. The medium was then replaced with 2 mL BSS containing either 10 μM 5-Ph-IAA or 0.1% DMSO. Embryos were transferred to the custom-made agarose chamber together with the treatment solution, oriented within the agarose pocket, and imaged using the spinning-disc confocal microscope.

### Bioinformatics

Structural models were predicted using AlphaFold 3.^53^ The protein sequences of Ol-RCC1 (XP_023820025) and Ol-Ran (XP_020562191) were retrieved from the NCBI database. The model with the highest predicted template modeling (pTM) score (0.86) and interface predicted template modeling (ipTM) score (0.84) was selected for structural analysis and visualized using ChimeraX. Nuclear localization sequences/signals (NLSs) were predicted using NLStradamus with a prediction cutoff of 0.5.^54^

### Image quantification

Line-scan analyses were performed using NIS-Elements software (version 5.41.00). Line widths are indicated in the corresponding figure legends, and mean fluorescence intensities are shown in the graphs. Cell size and mean nuclear RCC1 fluorescence intensity one frame before NEBD during the 4-cell division were measured using NIS-Elements. Graphs were generated using GraphPad Prism (version 10.3.0; Dotmatics).

### Illustration and movie preparation

Schematic diagrams were created using Adobe Illustrator 2025 (version 29.8.1; Adobe). Supplementary movies were prepared using NIS-Elements and Fiji.

### Statistics and Reproducibility

All statistical analyses were performed using GraphPad Prism (version 10.3.0; Dotmatics). Pairwise comparisons were performed using two-sided unpaired Welch’s *t*-tests, whereas comparisons among more than two groups were performed using Brown-Forsythe and Welch ANOVA followed by Dunnett’s multiple comparisons test. Statistical significance was defined at *p* < 0.05 and is indicated as \**p* < 0.05, \*\**p* < 0.01, \*\*\**p* < 0.001, \*\*\*\**p* < 0.0001; ns indicates not significant. No statistical method was used to predetermine sample size. Sample sizes were selected based on previous studies and were sufficient for the statistical analyses performed. Healthy fertilized eggs were randomly selected. Investigators were not blinded during data collection or analysis.

**Table 1:** Plasmids used in this study.

| No. | Name | Description | Related Figures | Source |
| --- | --- | --- | --- | --- |
| 1 | pTK1050 | pCS2+OsTIR1(F74G)-P2A-mCh-H2B | Fig. 2E-J; Fig. S2B, D; Fig. 3C | 18 |
| 2 | pTK1026 | pCS2+OsTIR1(F74G)-P2A-mCh-α-tubulin | Fig. S2C | 18 |
| 3 | pTK1073 | pCS2+mCh-α-tubulin | Fig. S2C | 18 |
| 4 | pTK1076 | pCS2+mCh-H2B | Fig. 3C; Fig. 5C, H, J; Fig. S4A | 18 |
| 5 | pYM7 | pCS2+OsTIR1(F74G)-P2A-Ol-RCC1-mCh | Fig. 3D, H | This study |
| 6 | pYM22 | pCS2+OsTIR1(F74G)-P2A- Ol-RCC1<sup>TM</sup>-mCh | Fig. 3D, F, H | This study |
| 7 | pYM33 | pCS2+Ol-RCC1<sup>TM</sup>-mCh | Fig. 3F; Fig. 5A, C, H, J | This study |
| 8 | pYM8 | pCS2+OsTIR1(F74G)-P2A-Ol-RCC1<sup>Δ45</sup>-mCh | Fig. 3F | This study |
| 9 | pYM31 | pCS2+Ol-RCC1<sup>Δ45</sup>-mCh | Fig. 3F; Fig. S5A | This study |
| 10 | pYM9 | pCS2+OsTIR1(F74G)-P2A-Ol-RCC1<sup>D207A</sup>-mCh | Fig. 3F | This study |
| 11 | pYM32 | pCS2+Ol-RCC1<sup>D207A</sup>-mCh | Fig. 3F; Fig. S5A | This study |
| 12 | pTK1044 | pCS2+mCh-Ol-HURP | Fig. 4A, B, F, H; Fig. S4B, D, E | This study |
| 13 | pTK1066 | pCS2+OsTIR1(F74G)-P2A-mCh-Ol-HURP | Fig. 4H; Fig. S4D-E | This study |
| 14 | pTK1042 | pCS2+mCh-Ol-KIFC1 | Fig. S4A | 27 |
| 15 | pYM6 | pCS2+Ol-RCC1-mCh | Fig. 5A, C, F, H, J | This study |

**Table 2:**
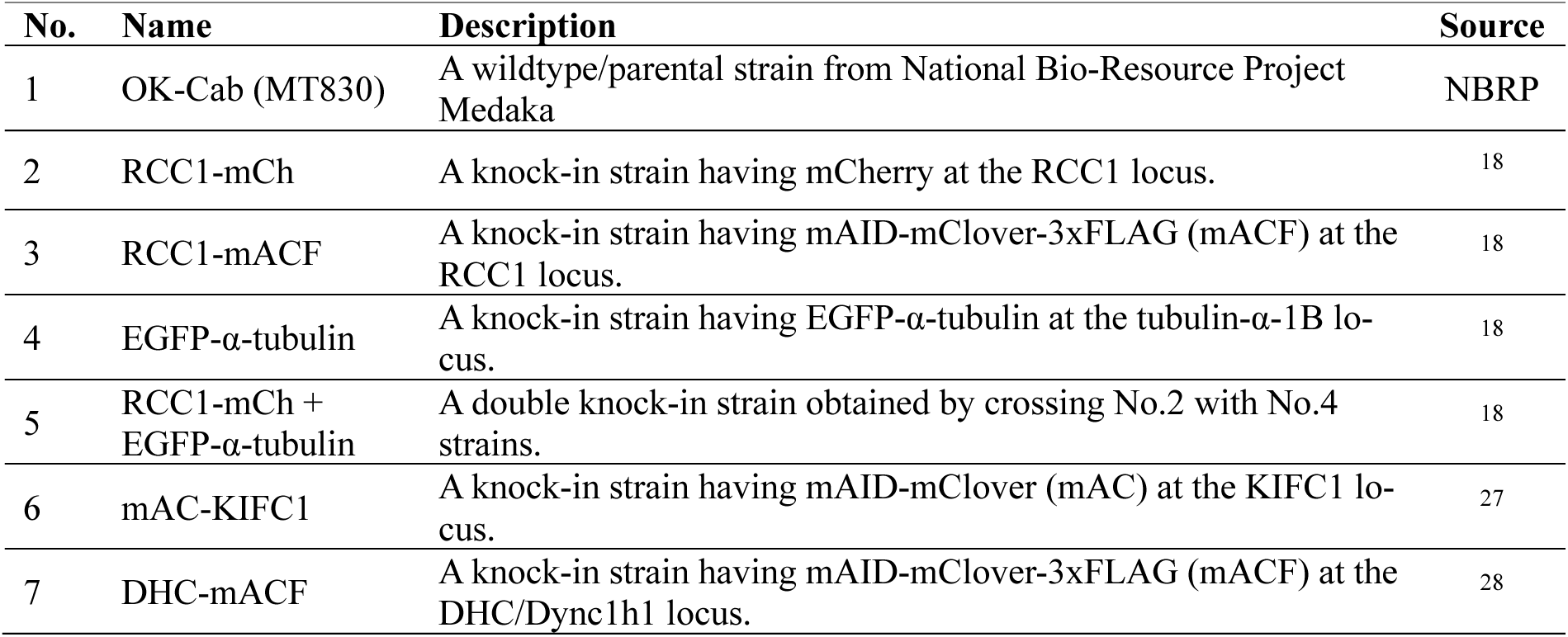
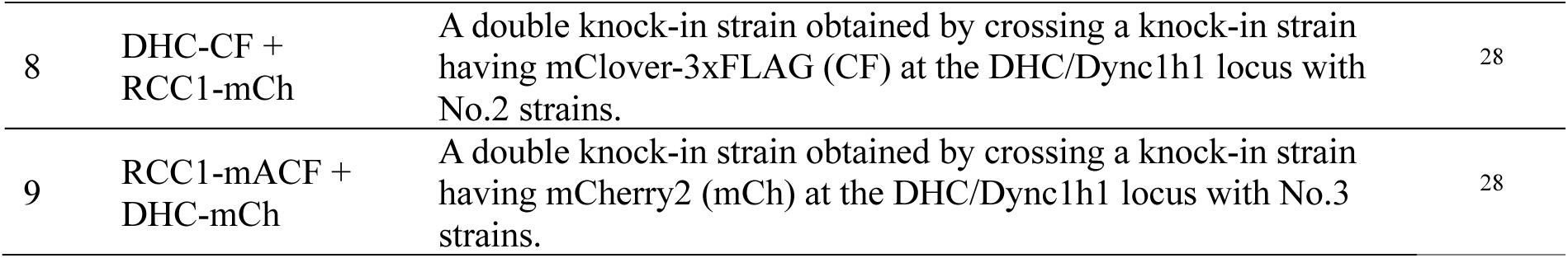
Medaka strains used in this study.

---

**Movie S1.** Related to Fig. 1D. Time-lapse movie of MIP images showing maternal RCC1 (RCC1-mCh, magenta) and paternal RCC1 (RCC1-mACF, green) in a fertilized medaka embryo during early embryogenesis.

**Movie S2.** Related to Fig. 3D. Time-lapse movie of MIP images showing chromosome segregation defects after endogenous RCC1 (RCC1-mACF, green) replacement with RCC1^TM^-mCh (magenta).

**Movie S3.** Related to Fig. 3F. Time-lapse movie of MIP images showing normal chromosome segregation after endogenous RCC1 (RCC1-mACF, green) replacement with RCC1^Δ45^-mCh (magenta).

**Movie S4.** Related to Fig. 3F. Time-lapse movie of MIP images showing normal chromosome segregation after endogenous RCC1 (RCC1-mACF, green) replacement with RCC1^D207A^-mCh (magenta).

**Movie S5.** Related to Fig. 4F. Time-lapse movie of a single z section showing mCh-HURP (magenta) and dynein (DHC-mACF, green) during spindle assembly in a two-cell-stage blastomere.

**Figure S1.**
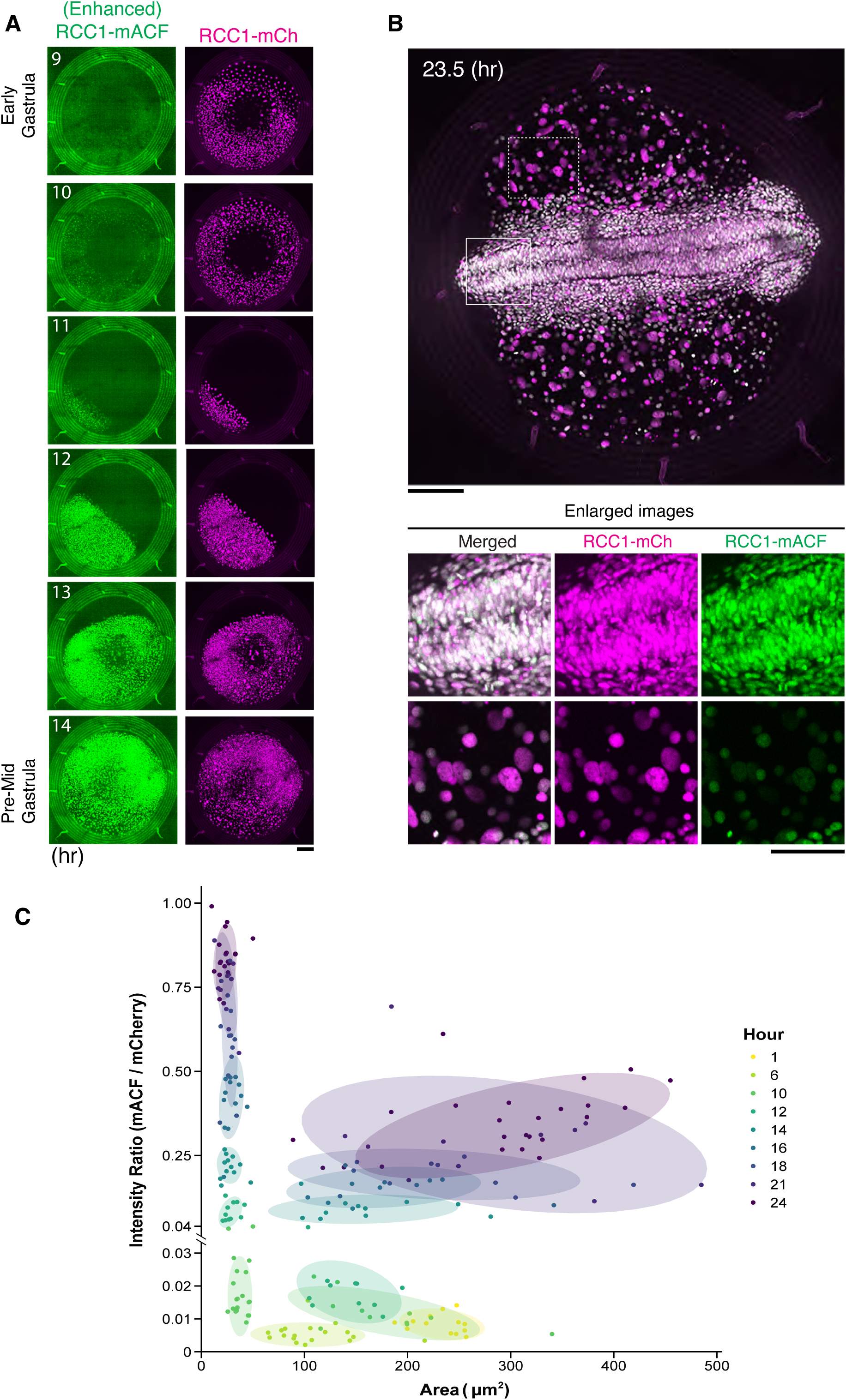
Characterization of maternal and paternal RCC1 during early embryogenesis, related to Figure 1. (A) Representative live images of maternal (RCC1-mCh) and paternal (RCC1-mACF) RCC1 from the early gastrula to pre-mid gastrula stages. Maximum-intensity projections (MIPs) of seven z sections are shown. (B) Enlarged view of the 23.5-hr time point from Figure 1D, showing comparable maternal and paternal RCC1 expression in embryonic cells (outlined by the solid box), whereas maternal RCC1-mCh was preferentially enriched in the larger nuclei of extraembryonic yolk syncytial layer (YSL) cells (outlined by the dashed box). These boxed regions are enlarged and shown below. (C) Nuclear area plotted against the paternal-to-maternal RCC1 fluorescence intensity ratio for individual nuclei. Colors indicate the imaging time points. Scale bars, 100 μm (A and B, top) and 50 μm (B, bottom).

**Figure S2.**
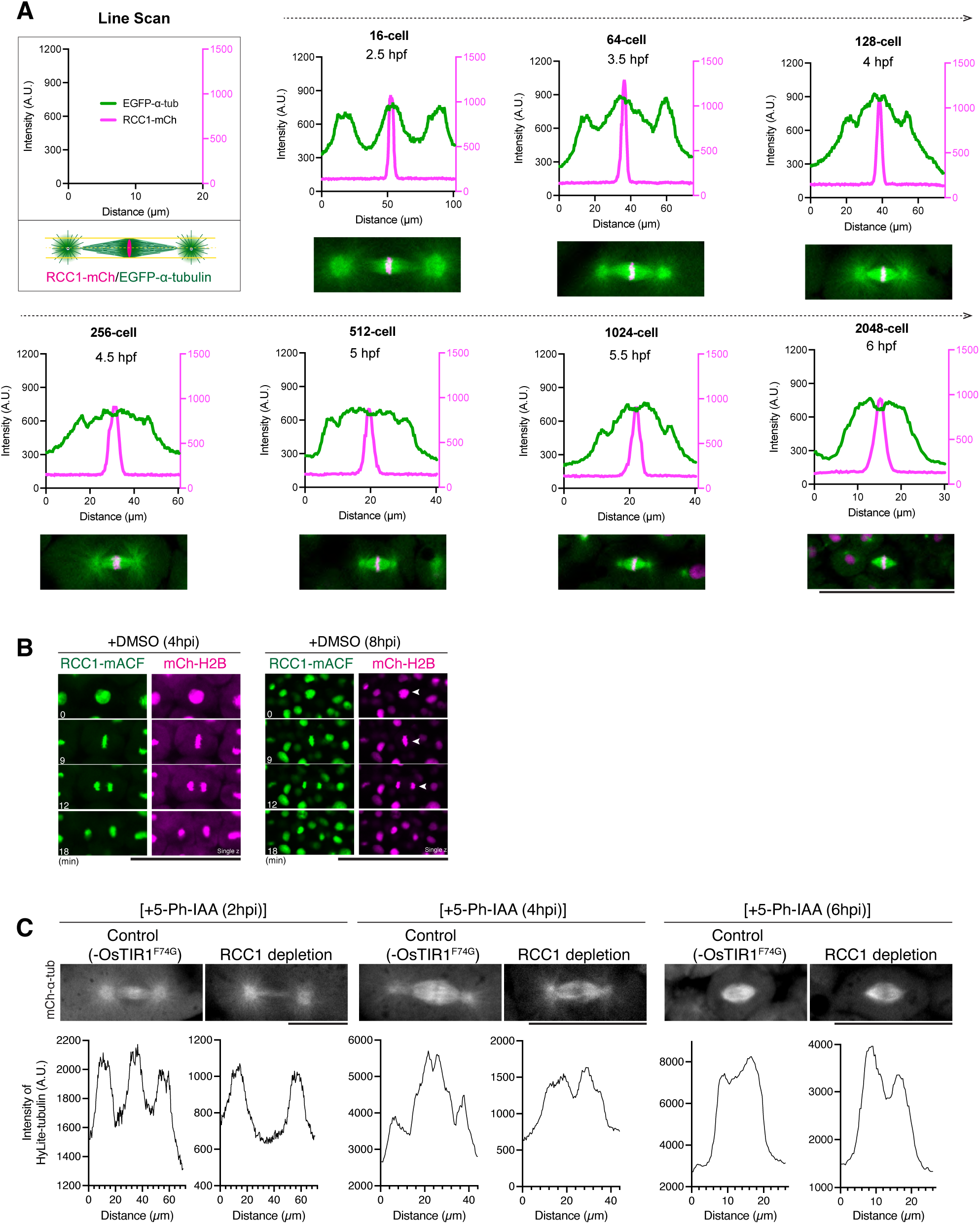
RCC1 requirement for embryonic spindle assembly changes during spindle remodeling around the early blastula stage, related to Figure 2. (A) Line-scan profiles (line width 8 px) of EGFP-á-tubulin and RCC1-mCh fluorescence intensity along the spindle axis in the representative live images shown below. (B) Representative live images of RCC1-mACF and mCh-H2B in embryos treated with DMSO at 4 or 8 hpi. (C) Representative live images (top) of mCh-á-tubulin in embryos with or without RCC1 depletion initiated at 2, 4, or 6 hpi, with corresponding line-scan analyses (bottom) along the spindle axis (line width 6 px). Scale bars, 100 μm (A, B) and 50 μm (C).

**Figure S3.**
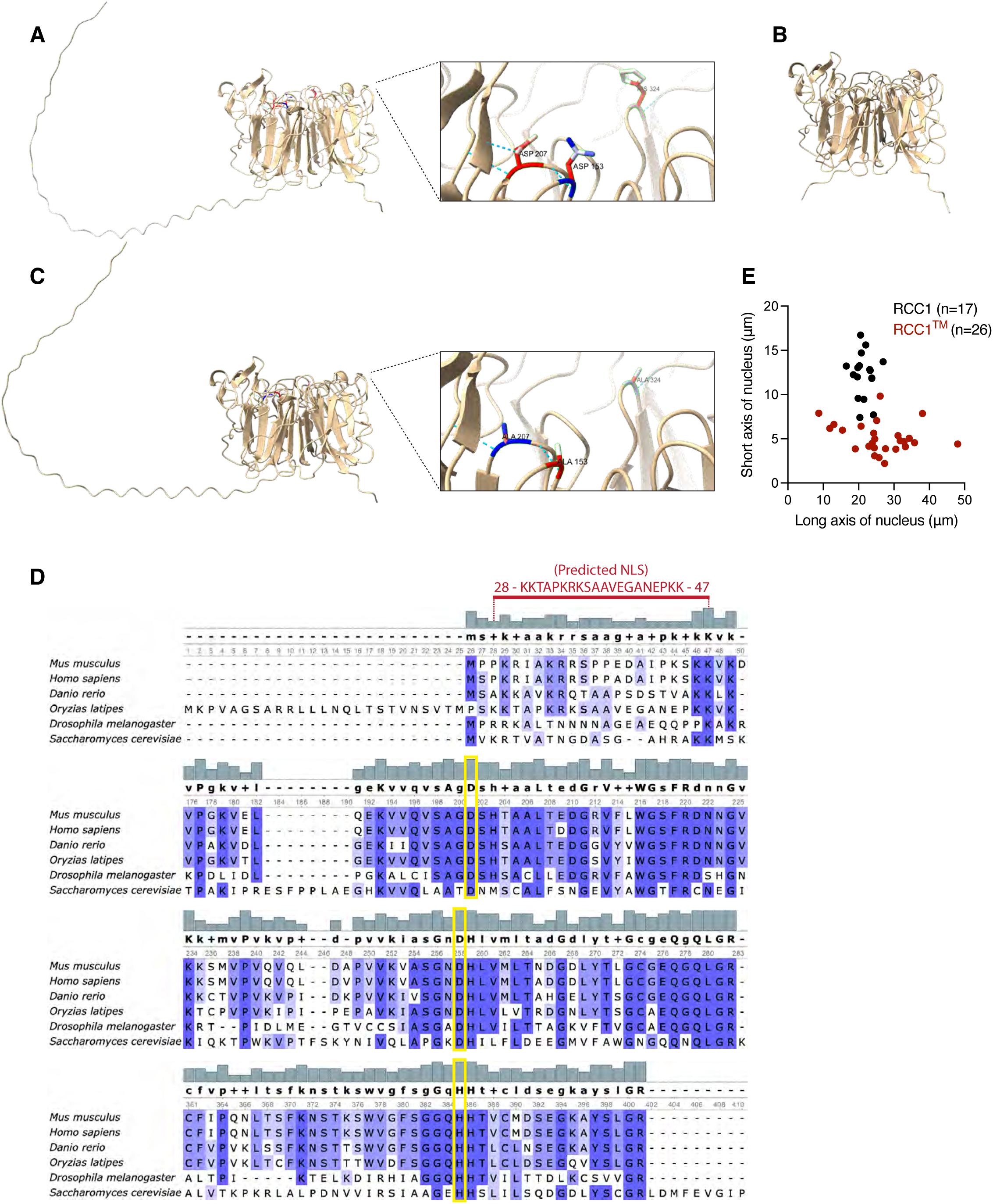
Structural features and conservation of medaka RCC1, related to Figure 3. (A) AlphaFold 3-predicted structure of medaka RCC1 (Ol-RCC1). (B) AlphaFold 3-predicted structure of Ol-RCC1^Δ45^. (C) AlphaFold 3-predicted structure of Ol-RCC1^TM^. (D) Amino acid sequence alignment of RCC1 proteins from *M. musculus* (NP_001184011), *H. sapiens* (NP_001260), *D. rerio* (NP_998343), *O. latipes* (XP_023820025), *D. melanogaster* (NP_523943), and *S. cerevisiae* (NP_011418), generated using Unipro UGENE. The alignment shows strong conservation of the GEF activity-related residues D153, D207, and H324 in medaka RCC1 (yellow boxes). The predicted NLS, identified using NLStradamus, is highlighted in red. (E) Scatter plots of nuclear size in RCC1-depleted 4-cell blastomeres after replacement with wild-type RCC1 or RCC1^TM^.

**Figure S4.**
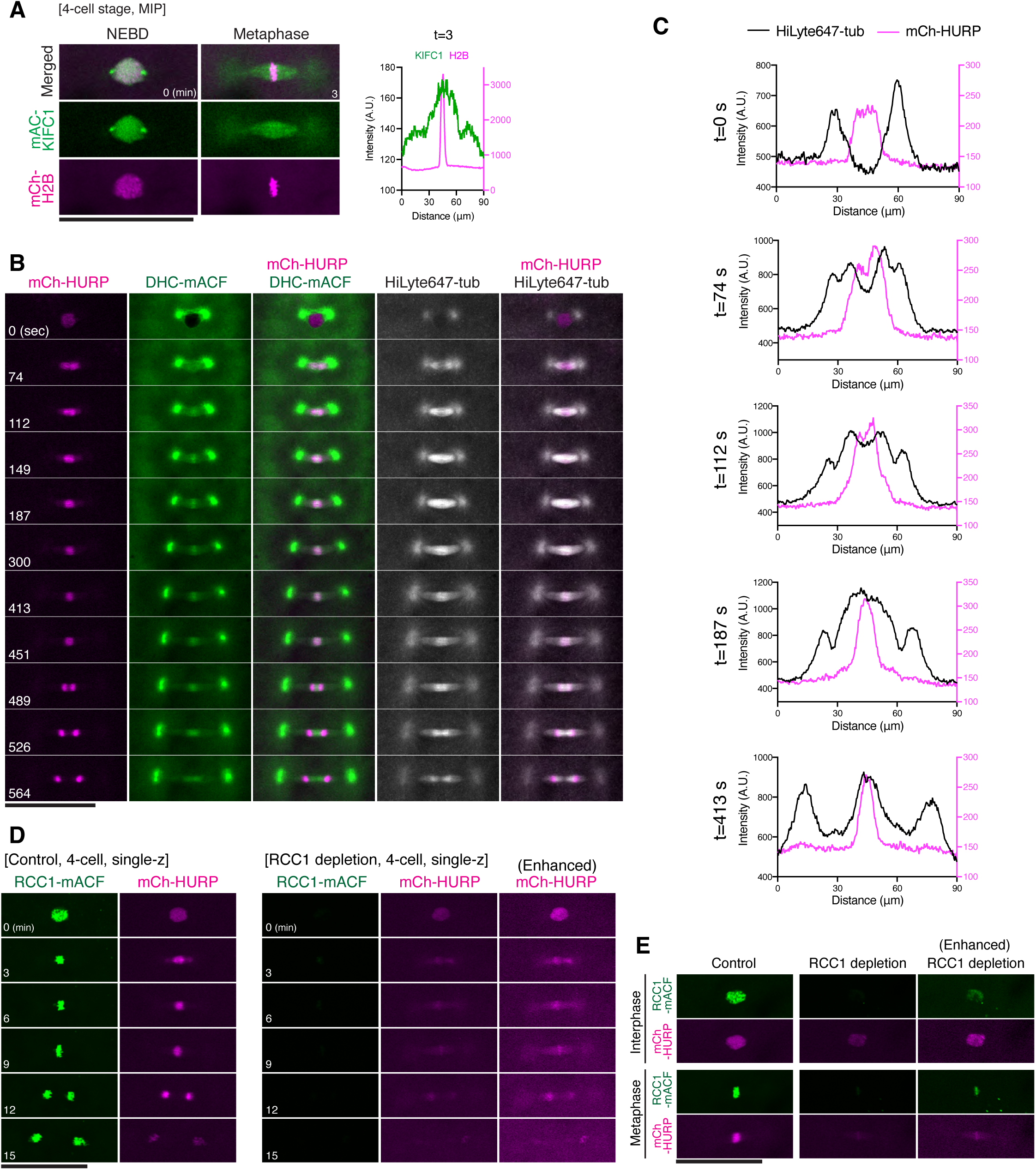
Spatiotemporal dynamics of HURP, KIFC1, and dynein during embryonic spindle assembly in the presence or absence of RCC1, related to Figure 4. (A) Representative live images of endogenous RCC1-mACF and exogenously expressed mCh-KIFC1, showing accumulation of KIFC1 around the metaphase spindle midplane in control embryos (left), but not in RCC1-depleted blastomeres (right). Corresponding line-scan analyses (line width 8 px) of KIFC1 and H2B fluorescence intensity across the metaphase spindle are shown on the right. (B) Representative live images of mCh-HURP and DHC-mACF together with HiLyte647-tubulin, showing the distinct patterns of spindle-midplane accumulation by mCh-HURP and DHC-mACF. HURP preferentially accumulates on spindle microtubules near chromosomes after NEBD (t=74). Enlarged images are shown in Figure 4F. (C) Corresponding line-scan analyses (line width 16 px) showing fluorescence intensities of HiLyte647-tubulin and mCh-HURP across the metaphase spindles in (B). (D) Representative live images showing disrupted localization of mCh-HURP around mitotic chromosomes during cleavage divisions in RCC1-depleted 4-cell blastomeres. Interphase (t=0) and metaphase (t=9) images are shown in Figure 4H. (E) Additional example showing reduced HURP accumulation at the spindle midplane in RCC1-depleted 4-cell blastomeres. Scale bars, 100 μm.

**Figure S5:**
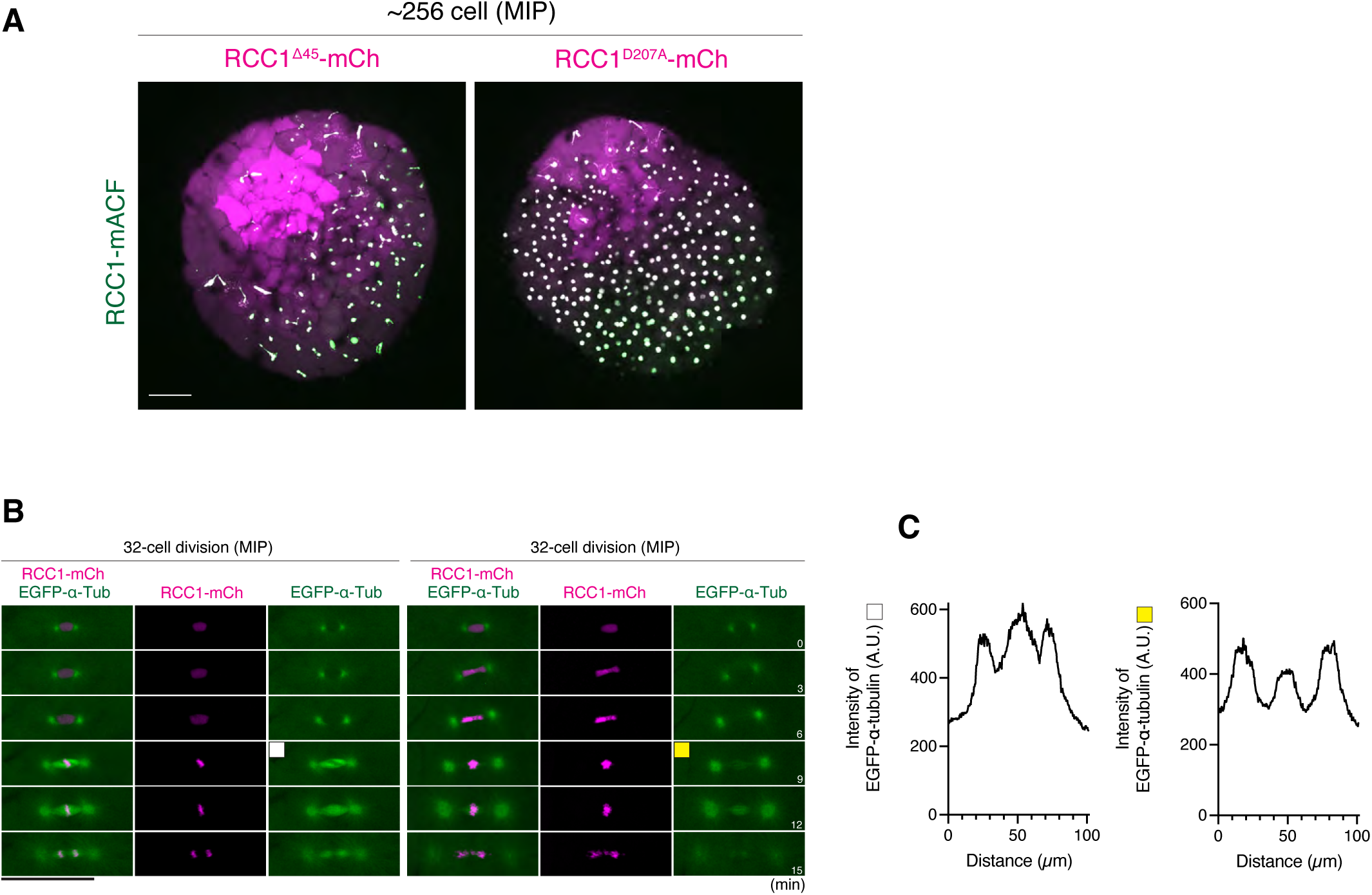
Overexpression of RCC1, RCC1^Δ45^, or RCC1^D207A^ disrupts spindle midplane organization, related to Figure 5. (A) Representative live images showing abnormal chromosome segregation following overexpression of RCC1^Δ45^ (left) or RCC1^D207A^ (right) at the 256-cell stage. (B) Representative live images of metaphase spindle midplanes in blastomeres expressing moderate (left) or high (right) levels of wild type RCC1 in 32-cell-stage embryos. (C) Line-scan analysis (line width 8 px) of EGFP-á-tubulin fluorescence intensity across metaphase spindles in (B). Scale bars, 100 μm.

